# Hierarchical Bayesian inference for concurrent model fitting and comparison for group studies

**DOI:** 10.1101/393561

**Authors:** Payam Piray, Amir Dezfouli, Tom Heskes, Michael J. Frank, Nathaniel D. Daw

## Abstract

Computational modeling plays an important role in modern neuroscience research. Much previous research has relied on statistical methods, separately, to address two problems that are actually interdependent. First, given a particular computational model, Bayesian hierarchical techniques have been used to estimate individual variation in parameters over a population of subjects, leveraging their population-level distributions. Second, candidate models are themselves compared, and individual variation in the expressed model estimated, according to the fits of the models to each subject. The interdependence between these two problems arises because the relevant population for estimating parameters of a model depends on which other subjects express the model. Here, we propose a hierarchical Bayesian inference (HBI) framework for concurrent model comparison, parameter estimation and inference at the population level, combining previous approaches. We show that this framework has important advantages for both parameter estimation and model comparison theoretically and experimentally. The parameters estimated by the HBI show smaller errors compared to other methods. Model comparison by HBI is robust against outliers and is not biased towards overly simplistic models. Furthermore, the fully Bayesian approach of HBI enables researchers to quantify uncertainty in group parameter estimates, for each candidate model separately, and to perform statistical tests on parameters of a population.

## Introduction

Across different areas of neuroscience, researchers increasingly employ computational models for experimental data analysis. For example, decision neuroscientists use reinforcement learning and economic models of choice to analyze behavioral and brain imaging data in reward learning and decision-making tasks [1,2]. The field of computational psychiatry uses these models to characterize patients and people at the risk of brain disorders [3–6]. Neuroimaging studies use models of neural interaction, such as dynamic causal modeling [7,8], as well as abstract models to analyze brain signals [1,9]. The success of these efforts heavily depends on statistical methods making inference about validity and robustness of estimated parameters across individuals, as well as making inference on validity and generalizability of computational models. A key theoretical and practical issue has been capturing individual variation both in a model’s parameters and additionally in which of several candidate models a subject expresses, which may also vary from subject to subject.

Computational models usually rely on free parameters, for example learning rate in reinforcement learning models, which often capture quantities of scientific interest but typically vary across individuals and must be estimated from data. A dataset includes a number of subjects, and often the question of interest is to characterize parameters in a population: Is choice consistency altered in patients with attention-deficit hyperactive disorders? Do cognitive enhancers, such as Ritalin, enhance learning rate at the population level? These questions are most naturally framed in terms of hierarchical models, which characterize both the population distributions over a model’s parameters, and also each individual subject’s parameters given the population distribution. Since these two levels are mutually interrelated, they are often estimated simultaneously, using methods like expectation maximization or sampling (MCMC). For example, the hierarchical parameter estimation (HPE) procedure [10,11] regularizes individual estimates according to group statistics, producing better individual estimates and permitting reliable group-level tests. Because subjects typically share underlying structure, hierarchical Bayesian approaches can leverage this this structure to yield better individual estimates, and to provide better predictions for unseen data, compared to approaches that fit each subject separately [12].

A second, and seemingly logically prior, question is which of several candidate models provides the best explanation for the data. This is important both for providing the setting within which to do parameter estimation, and also for investigating questions of scientific interest. Are rodents’ reaction times best explained by independent or competing accumulators? Do compulsive gamblers rely more on model-free reinforcement learning compared to controls? Importantly, in principle (and apparently in practice) the model expressed might also vary from subject to subject; thus modern model comparison techniques rely on estimating which of several models obtains for each subject [13]. Estimating such variation is important since by assuming that the same model obtains across all individuals (treating model identity as fixed effect) inflates significance for model comparison and makes it sensitive to outliers [13]. To estimate this variation, in turn, depends on the likelihood of each subject’s data given each model (and, thus, on each subject’s parameters for each model).

Intuitively, evaluating whether a model is a good model for a subject’s data precedes estimation of its specific parameter values; and indeed previous research has used separate tools to solve these two problems. But statistically, the two questions are actually interconnected, because individual parameters and hence individual fit depend on which subjects belong to the population that expresses the model. Here, we address this challenge from a fully Bayesian viewpoint. This work addresses issues of statistical inference over both parameters and models, which have remained elusive with the previous hierarchical methods.

Notably, although it is accepted (for the reasons discussed above) that the best-fitting model may vary from subject to subject, hierarchical parameter estimation (conducted separately) has typically assumed that the given model is expressed over all subjects, i.e. that it is a fixed effect. (And if multiple models are compared, these are each fit to the entire population.) This assumption biases parameter estimation, at both individual and group levels, because it entails that the estimated parameters for each individual subject *equally* affect group-level estimates, even though some members of the population may be better understood as expressing altogether different models. This same bias, in turn, affects the estimation of which subjects are best fit by each model.

In this work, we introduce a hierarchical and Bayesian inference method, which solves these problems by addressing both model fitting and model comparison within the same framework using variational techniques. Furthermore, our fully Bayesian approach enables us to assess uncertainty and provide a rigorous statistical test for making inference about parameters of a model at the population level, an issue that has been incompletely addressed in some previous hierarchical models. This paper is structured as follows. First, we highlight the main theoretical advances of our approach. A full formal treatment is given in the appendix. We then apply the proposed method to synthetic choice datasets as well as an empirical dataset to demonstrate its advantages over previous methods.

### Theory

Consider a typical computational modeling study in which data of a group of subjects have been measured and a set of candidate models are considered as possible underlying computational mechanisms generating those data. Such studies have generally two main goals: 1) to compare model evidence across competing models; 2) to estimate free parameters of models for each individual and their group-level distributions. All this is typically characterized in terms of inference in a hierarchically structured model of the data, which captures how each subject’s observations depend on their parameters, and the individual parameters on their group distribution.

The HPE procedure [10,11] employs a hierarchical approach to define the priors based on statistics of the group. This method typically assumes that for a particular model k, all individual parameters are normally distributed,

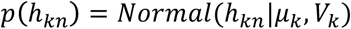

where *h*_*kn*_ is a vector of the free parameters of *k*th model for subject *n, μ*_*k*_ and *V*_*k*_ are the mean and variance parameters, respectively, indicating the prior distribution over *h*_*kn*_

HPE uses the expectation-maximization algorithm [14], a well-known iterative procedure, for obtaining estimating group parameters *μ*_*k*_ and *V*_*k*_ and individual parameters *h*_*kn*_ Every iteration of this algorithm alternates two steps: 1) an expectation step in which the individual parameters are estimated in light of the group-level distribution; and 2) a maximization step in which the group parameters, *μ*_*k*_ and *V*_*k*_, are updated given the current estimates of the individual parameters. Importantly, this update weights the individual subjects’ estimates equally; for instance, the update for *μ*_*k*_ is given by the average of subject level mean estimates (denoted *θ*_*kn*_) across all subjects:

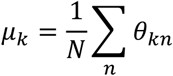

where *N* is the number of subjects.

Although HPE characterizes variation across subjects in the model parameters *h*_*kn*_ (that is, it treats those parameters as random effects), a critical assumption of the procedure is that the parameters for model *k* are estimated assuming that same model is responsible for generating data in all subjects. That is, the model identity is taken as a fixed effect, in contrast to random effects approach that assumes different models might be responsible for generating data in different subjects. The fixed effects assumption has two important implications: 1) for parameter estimation, group parameters, the group mean *μ*_*k*_ and variance *V*_*k*_, are influenced equally by all subjects, even those who would be better fit by some other candidate model j ≠ *k*; 2) for model comparison, the straightforward procedure (e.g. iBIC from [10,11]) is to compare models according to the sum of individual model evidences over all subjects, i.e. again treating the model identity as a fixed effect. Note that while it is possible to submit individual model evidence values (per subject and model) derived from HPE to a separate model comparison procedure that treats model identity as a random effect (such as random effects model selection [13]), these will be biased both from having been fit under the fixed effects assumption and also due to the optimization of the free group-level parameters. Altogether, violations of the fixed effects assumption can adversely influence both parameter estimation and model comparison.

Here, we extend HPE’s generative model with another level of the hierarchy, specifying for each subject which model generated their data. This is governed by a subject-specific multinomial random variable, itself drawn from a distribution controlling the proportion of each model in the population. This, in effect, merges the Bayesian model selection model from Stephan et al. [13] with HPE. We then lay out a procedure for joint inference over model identities and parameters, including quantifying the probability that each model is responsible for generating data for each subject. To achieve this goal, we take a full Bayesian approach in which the group parameters for each model, *μ*_*k*_ and *V*_*k*_, are also random variables. This also gives us a straightforward way to quantify the level of certainty in group-level estimations. We use variational Bayes [15,16], an extended version of expectation-maximization [17], which is able to deal with multiple latent variables in a probabilistic model. Since HBI is a variational framework, the resulting algorithm (Appendix A.3) is an iterative algorithm. On every iteration, the HBI performs 4 steps: calculates the summary statistics, updates its estimates of the posterior over group parameters, updates its estimate of the posterior over each individual parameter and finally updates its estimates of responsibility of each model in generating each individual data. The algorithm and other important mathematical issues are given in appendix A. Here, we highlight three main results. The mathematical proofs are given in appendix B.

As noted above, the HBI method estimates the probability of each subject’s dataset being generated by each model, or the *responsibility* of model *k* for generating data for subject *n, r*_*kn*_, which is expressed as (expected) probability. Larger values of *r*_*kn*_ (i.e. close to 1) indicate that model *k* is likely to be the true underlying model of the *n*th subject. In contrast, smaller values of *r*_*kn*_ (close to 0) indicate that model *k* is unlikely to be the underlying model for the *n*th subject. Based on the responsibilities, it is then possible to estimate number of subjects explained by each model, 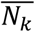

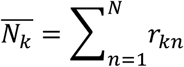

Thus 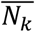 is always less than number of subjects and indexes the predominance of model *k* in the population. Furthermore, the fraction 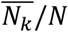 always lies between 0 and 1 and is a useful and intuitive metric for model comparison.

In practice, in many situations, researchers are interested in selecting a single “best” model (rather than relative comparisons among several) even in the face of variation in model identity across subjects. One way to accomplish this goal is to use 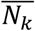 to compute the exceedance probability of each candidate model, a metric commonly used for model selection [13]. Exceedance probability is the probability that model *k* is more commonly expressed than any other model in the model space. Furthermore, the random effects approach enables us to quantify how likely the observed differences in model evidence is simply due to chance [18]. In this case, model selection is not statistically supported (although model comparison is valid). A metric called protected exceedance probability [18], which typically is more conservative than the exceedance probability, takes into account this possibility (see Appendix A.7). Altogether, the random effects approach results in a more robust model comparison and model selection, one less driven by outliers than fixed-effects methods. Note that previous attempts to do model selection at group level using exceedance probability assumed no hierarchy for parameter estimation, thus did not deal with the issue that parameter estimation was not properly conditionalized by group distributions based on model identity.

We noted above that an issue with the HPE is that the influence of subjects on the group parameters is equal. However, the comparable parameter in our approach, the mean of posterior distribution over *μ*_*k*_, denoted by *a*_*k*_, shows an important property: A subject’s effect on this parameter depends on the degree to which the model is estimated to be the underlying model for that subject. Specifically, this parameter, *a*_*k*_, is updated at each iteration as:

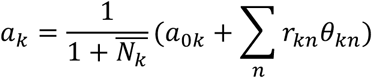

where *θ*_*kn*_ is the mean of the individual posterior and *a*_*0k*_ is the prior mean over *μ*_*k*_. The important point in this equation is that *a*_*k*_ is a *weighted* average of individual parameters, in which the weights are the corresponding responsibilities, *r*_*kn*_. This is not specific to the group mean, but it is rather a general feature of our approach: contribution of model *k* to group parameters is weighted according to the responsibility of model *k* in generating data in *n*th subject, *r*_*kn*_.

As mentioned above, another issue that has been incompletely treated in HPE is related to inference on parameters of a fitted model at the population level. Statistically, one needs the uncertainty of the estimated group mean, *μ*_*k*_, to be able to make inference on the corresponding parameter at the group level. Since parameters fitted by the hierarchical parameter estimation method are not independent but instead regularized according to the variance given by data, one cannot employ regular statistical tests, such as *t-*test, to test whether a specific model parameter is “significantly” different from zero. Using those tests on such parameters is biased in favor of generating a significant *p-*value (more false positives). The HBI framework solves this problem by quantifying uncertainty of the posterior over the group parameter. Specifically, it is possible to show that the posterior over the group parameter, *μ*_*k*_, takes the form of standard Student’s *t-*distribution centered at *a*_*k*_ with + 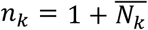 as degrees of freedom. The resulting *t*-value takes an intuitive form:

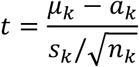

where *s*_*k*_ is the empirical deviance statistics. Therefore, 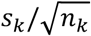 plays the role of standard error, which we call it *hierarchical error*. Note that the degrees of freedom of the test depend on the number of subjects (i.e. evidence) in favor of model *k* given by 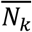, not the total number of subjects. Other group statistics, *a*_*k*_ and *s*_*k*_, are also weighted according to the responsibilities of model *k* in generating data of each subject (as formally obtained in Appendix A4).

## Results

In this section, we apply the proposed HBI method to synthetic and empirical datasets and compare its performance with that of HPE, as well as with Laplace approximation procedure, a non-hierarchical inference (NHI) method estimating parameters for each subject independently according to some fixed, *a priori* priors [19–22]. The HBI is general and could be applied to any type of data, such as choice data, reaction times, physiological signals and neural data. Since we are primarily interested in models of choice data, we focus on decision-making experiments. The details of simulations are given in the Supplementary Materials.

### Model comparison and parameter estimation

First, we simulated a dataset including 40 artificial datasets using two different learning models and a randomly generated reward sequence (binarized Gaussian random-walk). Both models maintain a value for each of the two possible actions and calculate a prediction error signal, δ, representing the difference between the seen reward and predicted value. On every trial, the action value gets updated according to the product of δ and a learning rate. The first model is a reinforcement learning model, in which the learning rate is a constant free parameter, α. The second model is a Kalman filter model in which the learning rate gradually decreases on every trial. The decreasing rate depends on a positive free parameter (representing observational noise), ω. Both models employ a softmax function together with an inverse-temperature parameter, β, to calculate the probability of each action according to corresponding expected values. Therefore, both models contain three free parameters and neither of them is nested within the other one.

The reinforcement learning and Kalman filter models were then used to simulate 10 and 30 artificial datasets, respectively. Parameters of these models were drawn randomly from normal distributions. Since parameters of these models have theoretical constrains, we used appropriate functions (sigmoid or exponential) to transform these randomly generated parameters. Using this procedure, we constructed a dataset of 40 artificial subjects, in which the true underlying model is known. We applied the HBI to this dataset to estimate parameters and model evidence given the sequence of actions. Simulations were repeated 20 times.

Figure 1 shows the results of the hierarchical Bayesian inference on this dataset. We first reported protected exceedance probability (Figure 1A), which represents the probability that each model is the most likely model across all subjects taking into account the null possibility that differences in model evidence is due to chance. This analysis revealed that the HBI has correctly identified the Kalman filter as the most likely model across the artificial datasets in all simulations with probability close to 1. Furthermore, HBI has indeed attributed about 10 and 30 artificial subjects to the reinforcement learning and Kalman filter models, respectively (Figure 1B). We then examined the performance of HBI in assigning the correct model at the individual level (Figure 1C). First, we found that the HBI has assigned the correct model to about 90% of all subjects (Figure 1C, inset). We then looked into the average of responsibilities for those artificial subjects whose underlying model was correctly assigned and for those cases whose model was erroneously assigned (Figure 1C). We found that the average of responsibilities estimated by HBI is about one for correctly identified cases and it is only slightly above chance for the rare cases that HBI failed to recognize the correct model.

**Figure 1.**
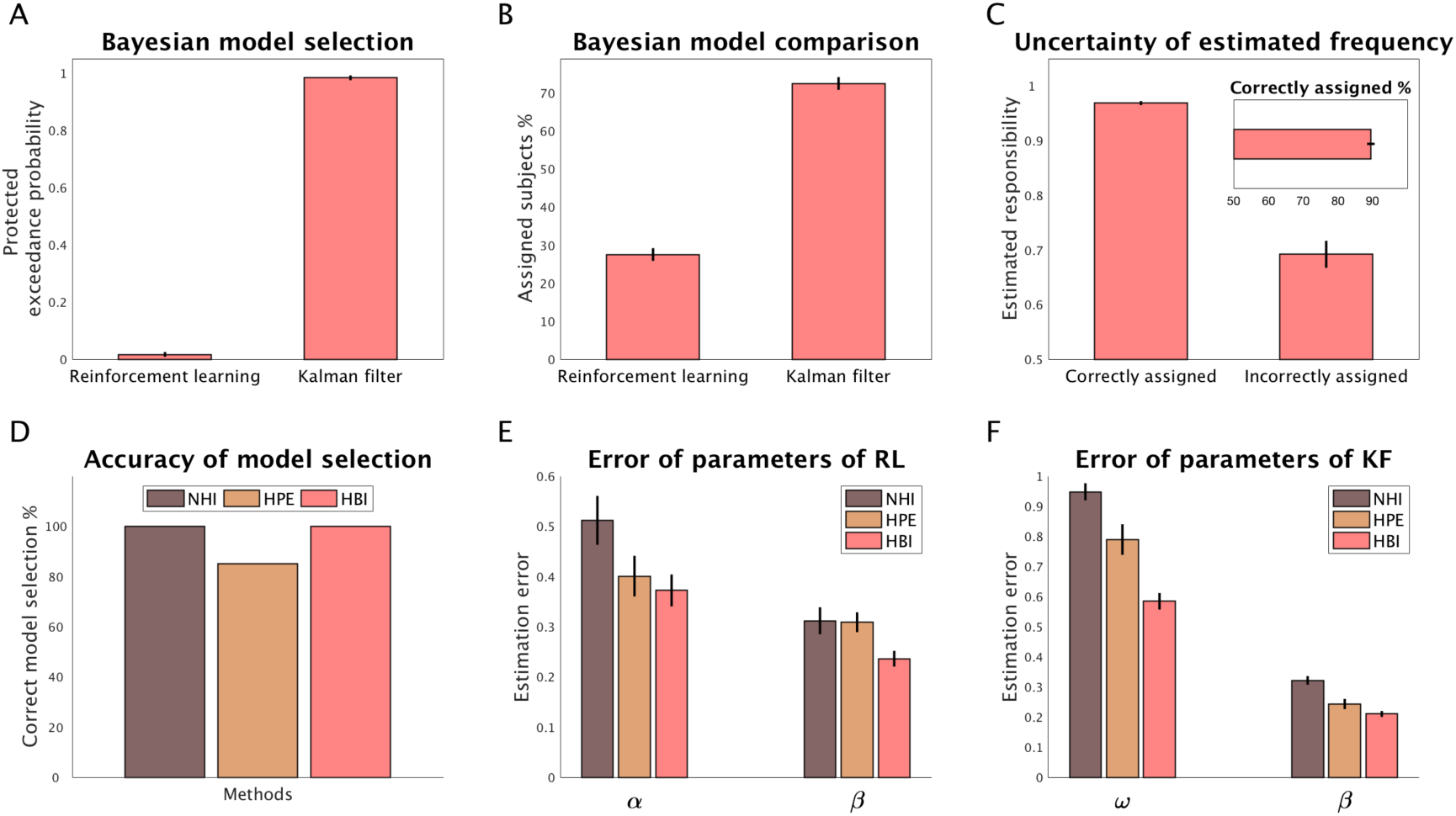
Performance of the HBI in a synthetic dataset. 10 and 30 artificial subjects were generated according to the reinforcement learning and Kalman filter models, respectively. A) Bayesian model selection using protected exceedance probabilities; B) Percentage of subjects explained by each model, estimated by the HBI. C) Uncertainty of HBI in estimation of responsibility for the correctly– and incorrectly– assigned subjects; Inset: percentage of correct assignment of the model by the HBI at the individual level. D) Comparison of accuracy of model selection with HPE and NHI; E, F) Error in estimating individual parameters of the reinforcement learning (E) and the Kalman filter model (F). Note that the errors are computed on the normally distributed parameters (not the transformed ones). The estimation error is defined as the absolute difference between estimated parameters and the true parameters. In all plots, error-bars are standard errors of the mean obtained across simulations 20 times. Abbreviations: HBI, hierarchical Bayesian inference; HPE, hierarchical parameter estimation; NHI, non-hierarchical inference; RL, reinforcement learning; KL, Kalman filter.

This means that the HBI method was quite certain when it was successful in identifying the true model and uncertain in cases in which it failed to recognize the true model.

We then compared performance of the HBI with the HPE method of Huys et al. [10,11] and NHI. In the latter, the model evidence across the group was quantified using random effects model comparison [13,18], which uses approximate individual evidence quantified by the Laplace approximation to compute group evidence. The model comparison of HPE is essentially fixed effects, in which evidence in favor of each model is equal to the sum of individual evidence measures quantified using local Laplace approximation and the penalty due to fitting group parameters [23] (see Supplementary Materials for details). In this set of simulations, all methods performed well in recognizing the most likely model (i.e. the Kalman filter) across all samples (Figure 1d), although the HPE performed worse than the other two models (failing 15% of simulations). In the next section, we examine limitations of HPE for model comparison more thoroughly.

We then investigated performance of these methods in parameter estimation. Estimation error, defined as the absolute difference between estimated parameters and true parameters used for generating data, was calculated. For both models and all parameters, the average error in parameter estimation by HBI was smaller than those by HPE and NHI (Figure 1E and 1F). Furthermore, HPE performed better than NHI in estimation across all parameters. These results were indeed theoretically expected. Unlike NHI, both HPE and HBI use group statistics to regularize parameter estimation for each individual. However, while HPE uses all subjects equally to regularize group parameters of a model, HBI weights individuals according to its belief that that model is responsible in generating each individual dataset. Further simulation analyses, in which the ratio of subjects expressing each model were different, confirmed these results (Supplementary Figure 1-2).

### Robustness of model comparison to outliers

We noted before that fixed effects model comparison using HPE is very sensitive to outliers. This is because fixed effects approaches sum up evidence across all subjects. If a few outlier subjects show large evidence in favor of a model, those usually impact model comparison adversely. In contrast, the HBI takes a random effects approach, in which the contribution of every subject in favor of each model is normalized according to the corresponding responsibility, which is a relative evidence measure with a maximum of one. In this section, we show a simulation analysis to demonstrate this point.

We took the same datasets generated in the previous simulations by the reinforcement learning and Kalman filter models. We then identified one outlier subject in that dataset that showed a large evidence in favor of the reinforcement learning model. This outlier subject was then used to create datasets with 1, 2 or 3 outliers by copying it 1, 2 or 3 times, respectively, and adding those copies to the original dataset.

We then compared the performance of NHI, HPE and HBI. Note that while NHI and HBI perform random effects model comparison, HPE perform a fixed effects model comparison. As shown in Figure 2, whereas the performance of HPE is very sensitive to outliers, the random effects model comparison of NHI and HBI are robust. Note that although NHI performs well in model selection here, we will demonstrate its limitations for model comparison in the next section.

**Figure 2.**
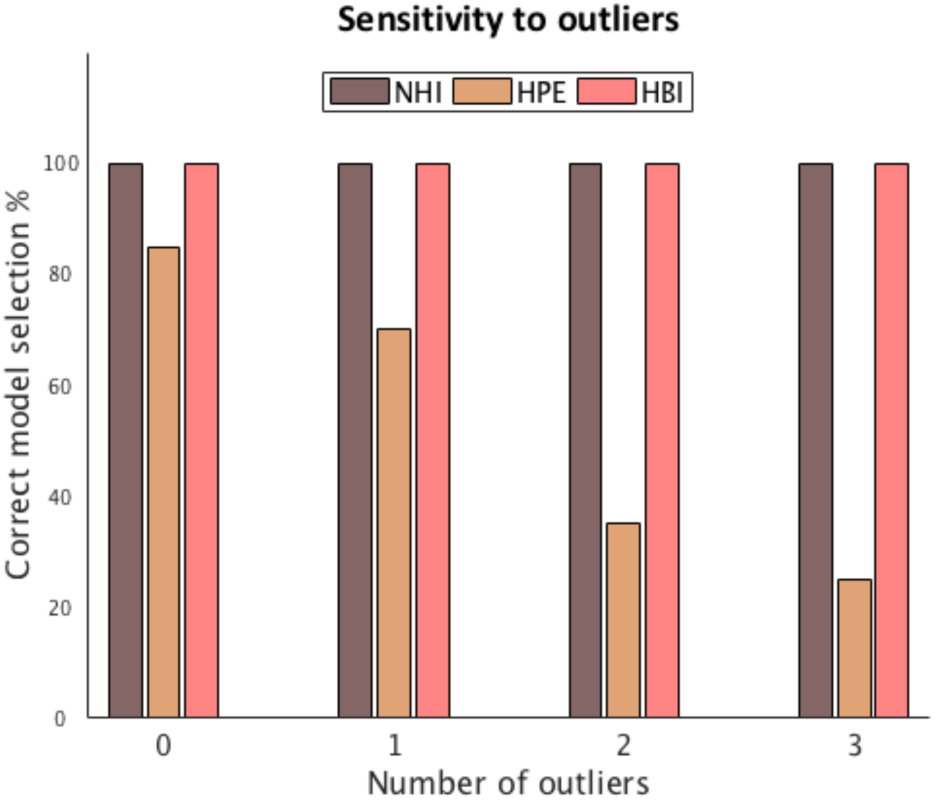
Sensitivity of model selection to outliers. The same 20 datasets simulated in the previous section were used as the base datasets (i.e. 0 outliers) and the effects of adding 1, 2 or 3 outliers to each dataset were examined. The HPE is severely sensitivity to outliers, while the other two (random effects) methods are robust.

### Model comparison and parameter estimation in nested models

We then considered a challenging problem in which the number of free parameters in two models is different and one model is a special case of the other one. Such problems are ubiquitous in studies using computational models and inference using hierarchical approaches is typically even more advantageous in this setting, as the variance explained by such models are more likely to overlap.

The first model was again assumed to be a reinforcement learning model with a constant learning rate parameter, α. The second model, however, was assumed to contain two different learning rates depending on whether the prediction error is positive or negative (dual-α reinforcement learning, commonly used to assess asymmetries in learning from positive vs negative prediction errors [24,25]). Both models use the same choice function, i.e., a softmax function with an inverse-temperature parameter, β. The reinforcement learning and the dual-α reinforcement learning models were then used to simulate 10 and 30 artificial datasets, respectively. Note that the reinforcement learning model is a nested case of the dual-α reinforcement learning, in which *a*^+^ = *a*^-^.

As Figure 3 shows, the HBI method was successful in model selection (i.e. recognizing the most likely model, Figure 3A) and attributed about 10 and 30 artificial subjects to the reinforcement learning and dual-α reinforcement learning models, respectively (Figure 3B). At the individual level, HBI assigned the correct model to each individual in 95% of all subjects and was also quite certain when it was successful in selecting the right model (Figure 3C). In contrast, in those rare cases in which HBI failed to recognize the correct underlying model, it assigned responsibility that was only slightly above chance.

**Figure 3.**
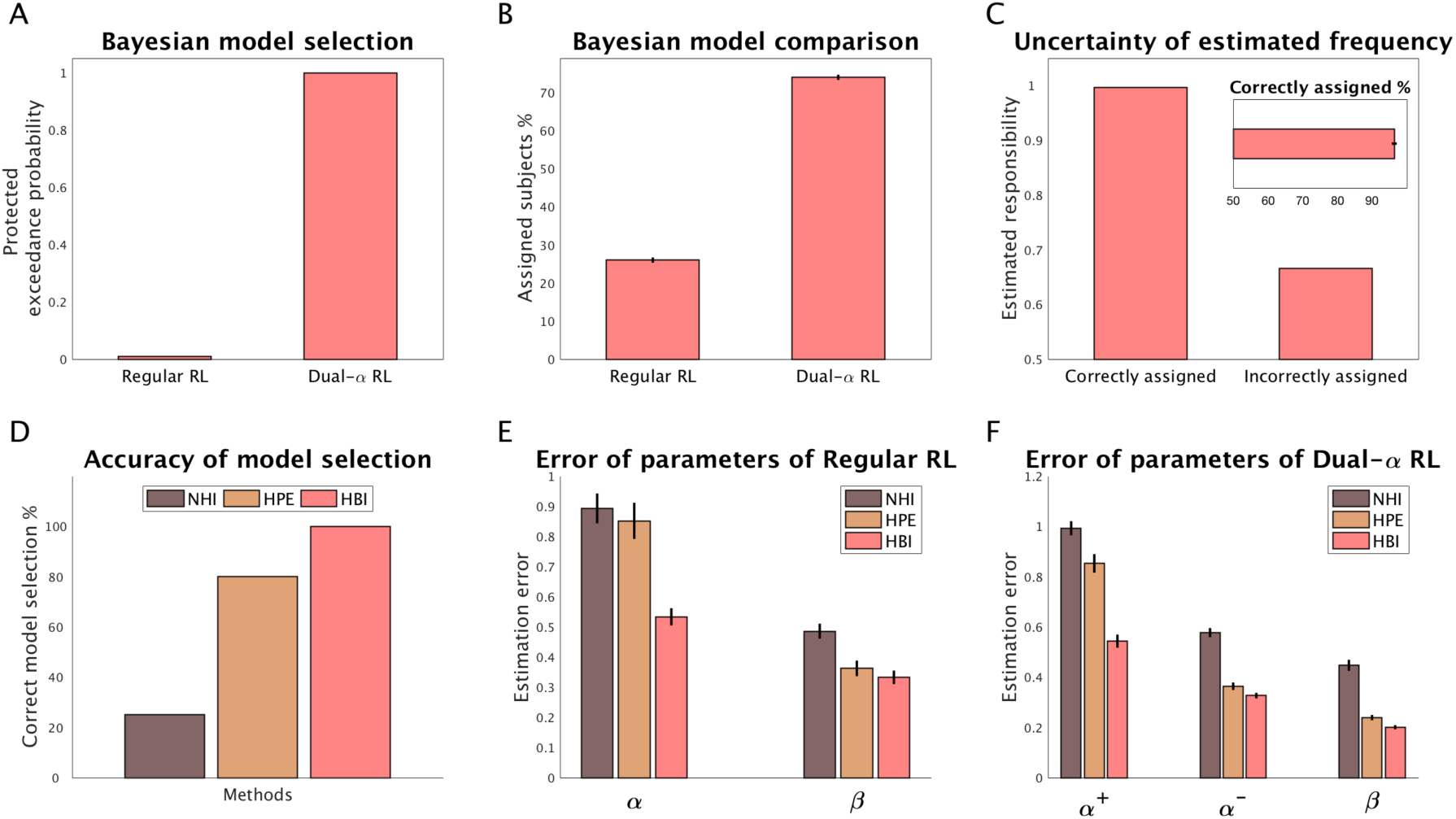
Performance of the HBI in a synthetic dataset. 10 and 30 artificial subjects were generated according to the reinforcement learning and dual-α reinforcement learning models, respectively. A) Bayesian model selection using protected exceedance probabilities; B) Percentage of subjects explained by each model, estimated by the HBI; C) Comparing uncertainty of HBI in estimation of responsibility for the correctly– and incorrectly– assigned subjects; D) Comparing accuracy of model selection by NHI, HPE, and HBI; E, F) Error in estimating individual parameters of the reinforcement learning (E) and the dual-α reinforcement learning model (F). Note that the normalized estimation errors are computed on the normally distributed parameters (not the transformed ones). The estimation error is defined as the absolute difference between estimated parameters and the true parameters. In all plots, error-bars are standard errors of the mean obtained across simulations 20 times. Abbreviations: HBI, hierarchical Bayesian inference; HPE, hierarchical parameter estimation; NHI, non-hierarchical inference; RL, reinforcement learning.

Next, we compared performance of the HBI with that of NHI and HPE. Here, NHI fails to choose correctly the most likely model in 75% of simulations. This is because non-hierarchical methods typically over-penalize more complex models, because they neglect the structure of the data. In particular, the issue is that a model with one additional parameter adds one independent free parameter per subject in the non-hierarchical case, which carries an excessive overfitting penalty, whereas these parameters are pooled by being drawn from a common distribution in the hierarchical setting, ensuring less overfitting and a more moderate complexity penalty. The HPE method performs much better, with a correct model selection in 80% of simulations. The HBI is successful in model selection in all simulations. We again examined performance of these methods in parameter estimation by calculating the absolute difference between estimated parameters and true parameters used for generating data. We found that errors in parameters estimated by HBI were smaller than those estimated by NHI or HPE for all parameters and both models, albeit the degree of improvement varies for different parameters. Further simulation analyses, in which the ratio of subjects expressing each model were different, confirmed these results (Supplementary Figure 3-4).

Note that the estimation errors of HBI are much smaller than those of HPE. Consider, for example, the learning rate parameter of the reinforcement learning model, α (Figure 3E). In generating the datasets for this analysis, α was assumed to be smaller than the learning rate parameters of the dual-α reinforcement learning model. Since the HPE uses average statistics across all subjects (even those generated by the dual-α model) to constrain parameters, the group average estimate of α by HPE was much larger than the true average. Therefore, the individual estimates of α by HPE are also tend to be larger than the true parameters, resulting in larger estimation error. The HBI does not have this problem because the group statistics are estimated using a weighted average, in which the weights are the corresponding responsibilities of models.

It is also important to note that all these methods are sensitive to amount of within-subject data (i.e. the number of trials). Importantly, the HBI is even more useful when there are limited number of trials (Figure 4). In this case, non-hierarchical methods, such as NHI, over-penalize complex models even more, as there are less data-points per subject to justify additional parameters. Furthermore, in this case, the HPE model selection performance is even more sensitive to outliers as it is more likely to have outliers when data per subject is limited (Figure 4A-C).

**Figure 4.**
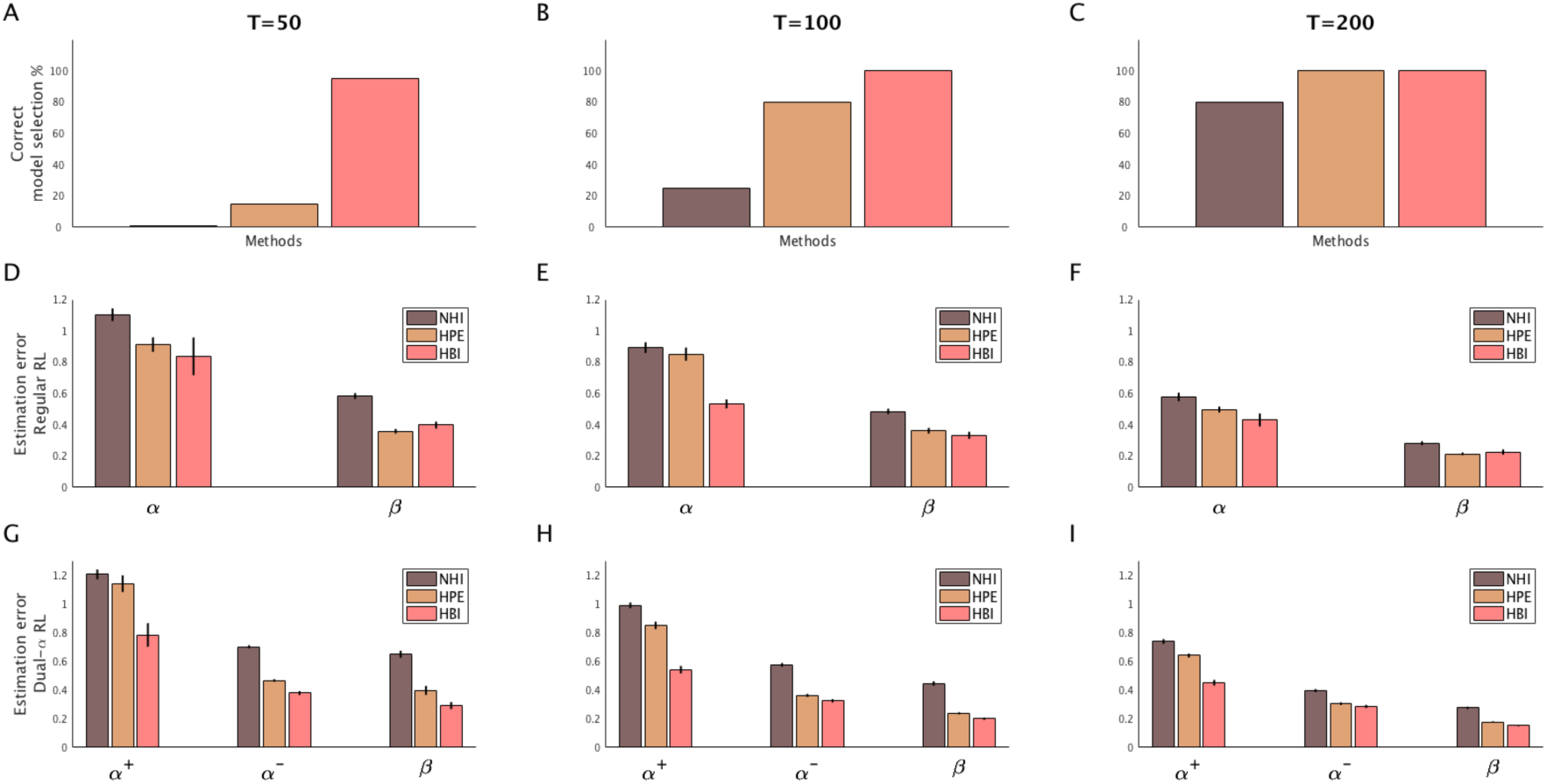
Performance of the HBI as a function of number of trials (T). 10 and 30 artificial subjects were generated according to the reinforcement learning and dual-α reinforcement learning models, respectively. These simulations were repeated 3 times with different number of trials per subject. A-C) Accuracy of model selection by NHI, HPE, and HBI for T=50, T=100 and T=200 trials; D-F) Error in estimating individual parameters of the reinforcement learning. G-I) Error in estimating individual parameters of the dual-α reinforcement learning. Note that the estimation errors are computed on the normally distributed parameters (not the transformed ones). The estimation error is defined as the absolute difference between estimated parameters and the true parameters. In all plots, error-bars are standard errors of the mean obtained across simulations 20 times. Abbreviations: HBI, hierarchical Bayesian inference; HPE, hierarchical parameter estimation; NHI, non-hierarchical inference; RL, reinforcement learning.

Note that hierarchical methods are also sensitive to amount of between-subject data (i.e. the number of subjects expressing each model). The difference between HPE and HBI is marginal in this case, although the greater benefit of HBI is when there are limited number of subjects as HPE is more likely to be impacted by outliers in those cases (Supplementary Figures 5). Furthermore, our simulations showed that when there is only one model in the model space, both HPE and HBI show (similar amount of) benefit compared with NHI in estimating parameters when there are less amount of within-subject data (Supplementary Figures 6), as reported in previous works [12].

**Figure 5.**
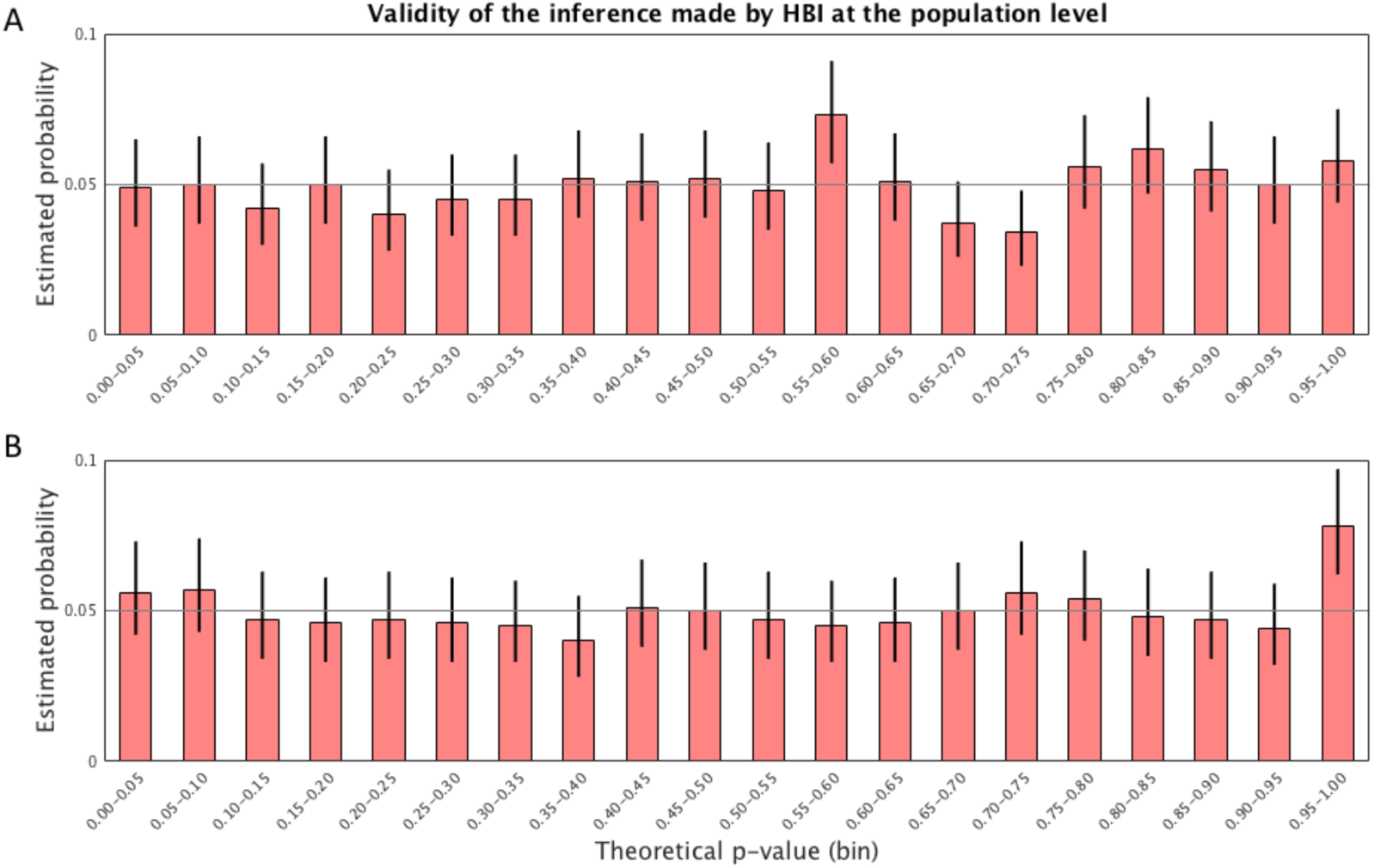
Performance of the HBI for statistical inference about model parameters at the population level. A bias parameter was generated under the null hypothesis at the group level in two simulations where there is only one model in the model-space (A) or there are more than one model in the model-space (B). Under the null-hypothesis, the resulting *p-*value of the group mean across all simulations is theoretically expected to have a uniform distribution. The distribution of estimated *p-*values by the HBI are plotted as a function of theoretical *p-*value in bins with the length of 0.05. The error-bars are 95% confidence intervals for the binomial distribution.

**Figure 6.**
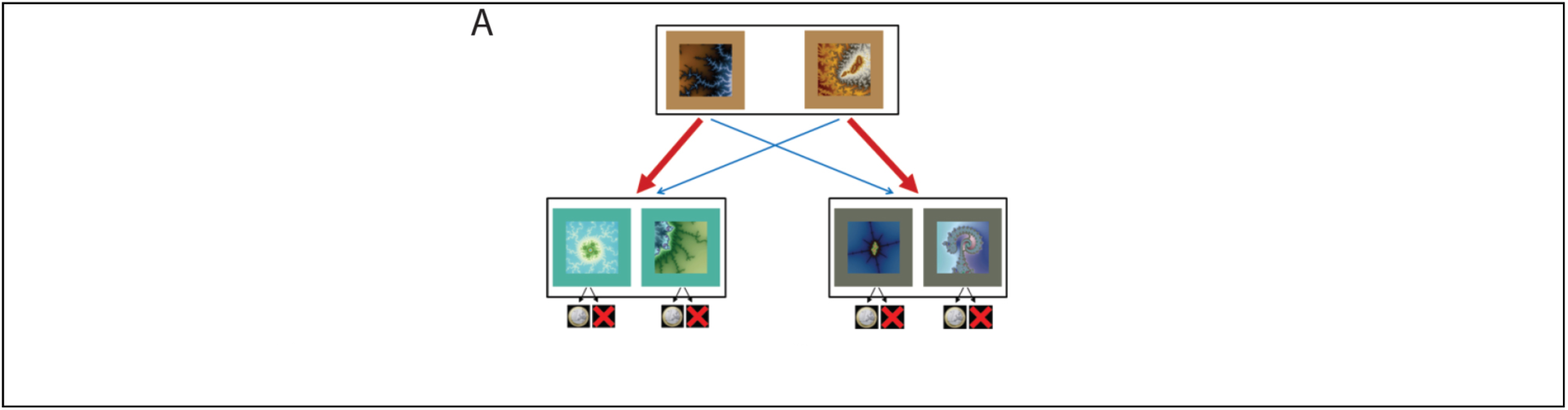
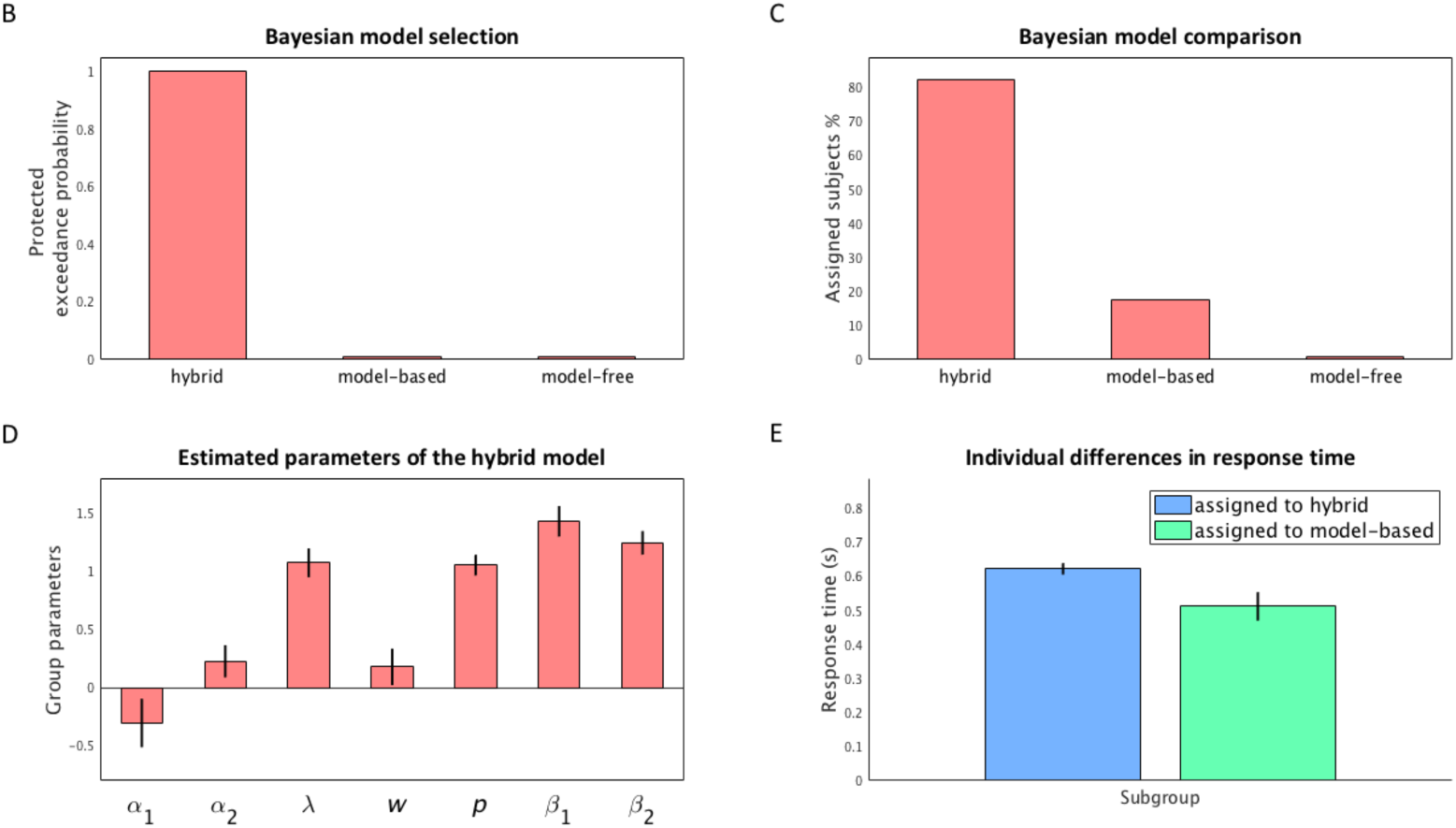
Performance of the HBI in an empirical dataset. A) the two-step Markov decision task [26]: every trial of the task comprises two steps. On each trial, subjects first made a choice between two fractal stimuli leading to one of the two different second-stage sets represented by different colors. Subjects then made another choice between two stimuli presented in the second-stage set. Each stimulus at the second-stage was associated with a specific probability of delivering a monetary reward. Critically, each choice at the first stage led predominantly (70%) to one of the two sets at the second stage (common transition), and less frequently (30%) to the other set (rare transition). This feature of the task allows for distinguishing contribution of model-based and model-free accounts on the first level choices. B) Bayesian model selection among the three models using HBI. Protected exceedance probabilities are plotted; C) Bayesian model comparison using HBI. Percentage of subjects explained by each model are plotted. HBI assigned almost 80% of subjects to the hybrid model. D) Estimated group mean parameters of the hyrbid model are plotted. Errorbars are corresponding hierarchical standard errors. The hybrid model [26] contains seven parameters: two learning rates, *α*_1_ and *α*_2_, for the first and second stage choice; an eligibility trace parameter, *λ*; a weight parameter, *w*, mixing model-based and model-free values (logit-transformed values plotted for all these parameters); a perserveration parameter *p*; and two decision noise parameters, *β*_1_ and *β*_2_(log-transformed values are plotted), for the first and second stage choices. E) Response time (mean and standard errors) for the first choice for the two subgroups that were assigned to hybrid or model-based account by HBI. Those who were assigned to hybrid showed slower response time.

### Inference about model parameters at the population level

We then focused on inference about parameters of a fitted model at the population level, and tested performance of the HBI using two simulation analyses. We focus on an example that represents a typical inference problem at the population level for parameters of a computational model.

Consider a situation in which subjects should learn stimulus-action-outcome contingencies. The subject’s task is to either to make a go-response by approaching the stimulus or to do nothing (i.e. no-go response). Furthermore, assume that the stimulus is either emotionally appetitive or aversive (e.g. a happy or an angry face cue), but the outcome value is independent of the emotional content of the stimulus. A question of interest is whether the emotional content (happy versus angry) of stimuli induces opposite biases in making a go response, regardless of action values (a form of Pavlovian to instrumental transfer). This is easy to test using a reinforcement learning model with one additional bias parameter, *b* (we call this model biased reinforcement learning). The bias is assumed to be *+b* for the emotionally appetitive stimulus and *–b* for the emotionally aversive stimulus. Thus, for larger values of *b*, the subject has a tendency to choose a go response after seeing the emotionally appetitive stimulus and a no-go response after seeing the emotionally aversive stimulus.

We simulated a dataset including 20 artificial subjects using this model and a randomly generated reward sequence (binarized Gaussian random-walk). Importantly, we assumed that the null hypothesis was true at the group level by drawing the bias parameter, *b*, randomly from a normal distribution with zero mean and variance of 1. A collection of 1000 datasets (each containing 20 artificial subjects) was simulated. The HBI method estimates the uncertainty of group mean parameters and corresponding degrees of freedom and gives the posterior belief about the group mean of parameters according to those estimates. One can then calculate the probability (*p-*value) that the estimated bias at the population level is generated by the null hypothesis. Note that in this example, all subjects are assigned to the biased reinforcement learning model as the model in the model space.

Under the null hypothesis, the HBI theory indicates that the *t*-statistics, 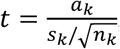, in which *a*_*k*_ is the posterior mean of group parameters, *s*_*k*_ is the empirical deviance statistics and 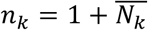, has a standard *t-*student distribution with *n*_*k*_ as degrees of freedom. Therefore, we expect that the corresponding *p-*value takes a uniform distribution between 0 and 1. This simulation analysis showed that the resulting *p-*value for the bias parameter indeed took a uniform distribution (Figure 5A).

Now we consider situations in which there is more than one model in the model space. Here, the HBI first infers the number of subjects explained by each model, 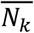, and then quantifies hierarchical errors and degrees of freedom according to 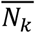 and the empirical group mean and deviance statistics. Therefore, we considered the same stimulus-action-outcome learning experiment as above and simulated a dataset including 40 artificial subjects. Data for half of subjects were generated using the same biased reinforcement learning model and data for the other half were generated using the dual-*α* reinforcement learning model explained in previous simulations. We again assumed that the null hypothesis is true at the group level by drawing the bias parameter randomly from normal distribution with zero mean and variance of 1. Again, a collection of 1000 datasets (each containing 40 artificial subjects) was simulated and the distribution of corresponding *p-*value was generated. Consistent with the theory, this simulation analysis showed that the resulting *p-*value for the bias parameter well follows a uniform distribution (Figure 5B).

### Empirical dataset: HBI reveals meaningful individual differences

We then applied the HBI method to empirical choice data from 31 subjects performing the two-step Markov decision task introduced by Daw et al. [26]. This task is a well-known paradigm to distinguish two behavioral modes, model-based and model-free learning [27–29]. Previous works have shown that there are important individual differences in this task [26,30], especially in the degree to which people employ a model-based strategy, and those differences are related to neuroanatomical [30,31], psychological [32], genetic [33] and psychiatric [34] trait scores. Daw et al. [26] have proposed three reinforcement learning accounts, a model-based, a model-free and their hybrid (which nests the other two and combines their estimates according to a weight parameter), to disentangle contribution of these two behavioral modes on choices. Here, we skip the details of the models and focus on application of the HBI to a model space consisting of model-free, model-based and hybrid accounts. The dataset used for this analysis have been reported elsewhere [30].

Before analyzing the empirical dataset, we did a simulation analysis of this task and model space. We verified that the HBI recovers the parameters of the models better than alternative methods. In particular, the critical weight parameter of the hybrid model, which determines the degree that each account influences behavior, was significantly better recovered by the HBI than the other methods (in all 20 simulations, HBI did better than both HPE and NHI, Supplementary Figure 7). We then applied the HBI on the empirical dataset of this task (Figure 6). As Figure 6B shows, consistent with previous findings on this task, the hybrid model accounts best for choices in this task with a protected exceedance probability close to 1. About 25 (%81 of all subjects) and 6 subjects have been assigned to the hybrid and model-based models, respectively, while the model-free took no responsibility (Figure 6C). The estimated group mean and the corresponding hierarchical errors of the hybrid model are plotted in Figure 6D. Table 1 shows the weighted average and standard deviation computed by the HBI for the hybrid model in which the weights are given by responsibilities taken by the hybrid model in explaining data of each individual subject.

We also performed further analysis testing whether individual differences found by the HBI generalize to individual differences in conceptually related, yet independent, data. We reasoned that subjects showing a hybrid strategy might be slower in their choice, as the hybrid model requires combining of model-based and model-free values (which in some trials might be in conflict). Therefore, we looked at the median of response time across all first-level choices for each subject and tested whether there is a difference in response times between those subjects who employed a hybrid strategy and those who employed a model-based strategy as estimated by the HBI. As Figure 6E shows, the subgroup attributed to the hybrid model by the HBI showed slower response time compared to those subjects attributed to the model-based account (p=0.03, Wilcoxon test). A similar analysis using individual differences found by the NHI revealed no significant difference in response times. These results suggest that HBI reveals meaningful individual differences generalizing to unseen data.

## Discussion

In this work, we have introduced a novel method, a hierarchical and Bayesian inference framework, for parameter estimation and model comparison. The HBI framework is hierarchical in the sense that parameters at the individual level are regularized by statistics across all individuals in the group. The HBI framework is Bayesian in the sense that all uncertainties at both individual and group levels are represented by probability distributions. The HBI framework has major theoretical advantages over current state-of-the-art methods, mainly because it combines two sorts of inference (about model identity and model parameters) in a single hierarchical model, which are interdependent but have previously been treated separately. Our simulation results demonstrated these advantages experimentally.

In this work, we took an empirical Bayes approach [35,36], in which priors are constructed based on data. In other words, parameters at the individual level are regularized by statistics across all individuals in the group. Furthermore, we took a so-called random effects approach to model identity [13], which indicates that different models might underlie data in different subjects. This is in contrast to previous hierarchical methods for model fitting, which assume the same model underlie data in all subjects (fixed effects assumption). The random effects approach to hierarchical inference has important consequences for both parameter estimation and model comparison. Moreover, we took a fully Bayesian approach by quantifying uncertainty at the group level, which enabled us to develop statistical tests about group parameters and to quantify corresponding statistical errors.

Empirical Bayes methods play an increasing role in modern statistics. These methods essentially take a hierarchical approach, by assuming that individual data are generated based on the probabilistic properties of population. This hierarchical approach has important consequences. The most important consequence is that they provide a promising solution to the classical problem of priors in Bayesian statistics by providing informative, yet objective, priors at the individual level. Furthermore, unlike non-hierarchical methods, model comparison based on these methods is not biased towards too simple models. This is because non-hierarchical methods assume that extra parameters of a complex model are independent. For example, consider a model space in which the more complex model has one extra free parameter and there are 40 subjects in the dataset. Fitting the dataset with the complex model using non-hierarchical methods introduces 40 additional independent free parameters, driving an excessive penalty for overfitting. The hierarchical approach, however, assumes that the individual parameters are dependent, as they are all generated according to the same distribution. Modeling this hierarchical dependency enables those methods to avoid penalizing complex models excessively. Our simulation results demonstrate this point experimentally (Figure 3D). While the non-hierarchical method failed to select the correct model with one additional parameter, the HBI was successful in selecting the correct model (Figure 3D).

The HBI method introduced in this paper is built based on the random effects view that different models might underlie data in different subjects. Taking this view enabled us to address problems caused by taking the model identity as a fixed effect in some hierarchical parameter estimation procedures. For parameter estimation, the fixed effects assumption biases the group parameters because it assumes that all subjects contribute equally to the group parameters. The proposed HBI framework solves this problem by weighting contribution of each subject to group statistics by the degree to which that model is likely to be the true underlying model for that subject (Figures 1 and 3). For model comparison, the fixed effects assumption leads to oversensitivity to outliers [13] as the evidence across the group is driven by the sum of individual evidences. Our simulation results (Figure 2) showed that only a few outliers change lead to incorrect model selection inference made by the fixed effects assumption. The proposed HBI method solves this problem by normalizing individual evidence across all candidate models. Specifically, the HBI framework quantifies the responsibility of each model *k* in generating each subject data, a metric lying between 0 and 1. For every subject, the responsibility sums up to 1 across all candidate models as it partitions probability space among those models (see [13,18] for a similar approach). It is then easy to compare models by enumerating responsibilities across the group in favor of each model or by estimating the most likely model.

Another major contribution of this paper is to provide a statistical solution to the inference problem at the group level using hierarchically fitted parameters. For models fitted by a non-hierarchical method, such as maximum likelihood or Laplace approximation, it is statistically valid to use conventional statistical tests on fitted parameters to make inference at the group level. However, for datasets fitted by a hierarchical method in which the individual fits are regularized according to statistics of the group data, conventional statistical tests are in some cases not valid, because the parameter estimates are non-independent from subject to subject. Our fully Bayesian approach enabled us to address this issue. This is because the HBI infers the posterior distribution of group parameters. Our method provides an intuitive solution to this problem in the form a *t-*statistic, in which all the group statistics are computed according to the estimated responsibilities of the corresponding model in generating each individual data. Thus, the HBI quantifies the uncertainty of the group parameters and thereby the corresponding hierarchical errors. Simulation analyses (Figure 5) highlighted this point experimentally by showing that *p-*values computed by the HBI under the null hypothesis (i.e. when the group parameter is normally distributed around zero) follow a uniform distribution. Therefore, the HBI framework enables researchers to make statistical claims about parameters at the group level.

In addition to model comparison, the HBI framework can also be used for model selection in situations where the goal is to select one of the models as the best model across the group. Exceedance probability is a metric proposed [13] to perform model selection using a random effects approach. An important revision of this metric called protected exceedance probability [18] also takes into account the possibility that none of the models in a model space is supported sufficiently by data, i.e. the differences in model evidence are due to chance. As the HBI framework treats model identity as a random effect, it is possible to compute exceedance and protected exceedance probabilities.

There are increasing efforts to exploit advances in computational modeling for understanding mental disorders [3–6]. Recent works, however, have started to tackle challenges related to quantifying uncertainty in diagnosis and also in evaluation of treatment effects. For example, hierarchical unsupervised generative modeling, have used Monte-Carlo and variational methods to identify cluster of subjects showing similar patterns of neural connectivity [37,38]. HBI also offers a promising solution by quantifying uncertainty in assigning models to data generated by a single case. Our simulation (Figure 1C and 2C) showed that HBI assigns probabilities that are close to chance in cases in which model identification goes wrong. This can help us to move towards better diagnosis and precise evaluation of different treatments [39].

In summary, the HBI framework proposed in this work rests on a hierarchical view of both hypothesis testing (i.e. model comparison) and parameter estimation for multi-subject studies and thus provides a generic framework for statistical inference. Moreover, the HBI framework runs fully automatically and it does not rely on hand tuning of parameters. Therefore, we expect this method to be useful for a wide range of studies testing different hypotheses in a multi-subject setting. This includes not only computational models of learning and decision making, but also any statistical models of brain or behavior.

## Appendix A

In this appendix, we give a formal treatment of the HBI framework. First, we give the probabilistic model underlying HBI. In A.2, our approach for making inference (the full proof is given in appendix B) and related assumptions are given. In A.3, the HBI algorithm is presented. In A.4, we show how the problem of statistical inference about group parameters is solved in HBI. In A.5, we show how HBI can be used for making inference about a new subject. Finally, in A.6, we highlight some practical points, such as the initialization of the parameters and their settings.

### A.1 Probabilistic model

We begin by describing the probabilistic model of the HBI. Consider an observed dataset **X** = {**x**_1_, …, **x**_*N*_} where **x**_*n*_ is the dataset (e.g. choices) of *n*th subject and *N* indicates the number of subjects and a model-space including *K* candidate models, *M*_1_ *… M*_*K*_. Moreover, suppose that the prior probability of each model in the population is given by **m** = {*m*_1_, *…, m*_*K*_}. For each dataset, **x**_*n*_, we assume that there is a latent variable **z**_*n*_ comprising a 1-of-K binary random vector, in which *z*_*kn*_ is one if **x**_*n*_ generated is by the *k*th model. Thus, the probability of the latent variable across all subjects, **Z** = {**z**_1_, …, **z**_*N*_}, is assumed to have a multinomial distribution,

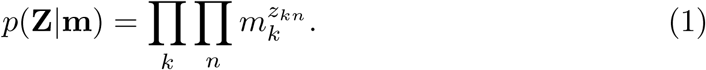

Each model *M*_*k*_ in the model-space is supposed to compute the probability of a given dataset (e.g. a set of choices) given a set of parameters, **h**_*kn*_ For example, the reinforcement learning model computes the probability of choices using two parameters: a learning rate and a decision noise parameter. The number of models and their structures depend on specific scientific questions. Here, we take a general approach by making no specific assumption about the number of models, *K*. Thus, the *k*th model in the model-space, *M*_*k*_, computes the probability of dataset **x**_*n*_ given the parameter vector **h**_*kn*_, which is denoted by *p*(**x**_*n*_|**h**_*kn*_, *M*_*k*_). Note that the number of parameters in model *k*, denoted by *D*_*k*_, might be different across models. Since data for each subject is generated by one of the models, which is denoted in the binary vector **z**_*n*_, the probability of the observed dataset given the model-space is

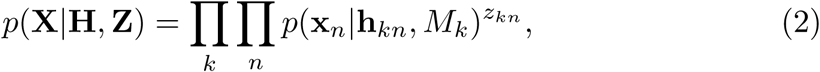

where **H** denotes all the parameters across all participants and models. The parameters of *k*th model are assumed to have a multivariate normal distribution with mean ***µ***_*k*_ and precision matrix **T**_*k*_,

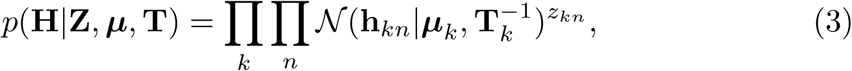

where **T**_*k*_ is a diagonal positive-definite matrix.

We also introduce a distribution over model frequencies, **m**. Since this is a probability over probabilities (which sum to one), we use the Dirichlet distribution as the prior:

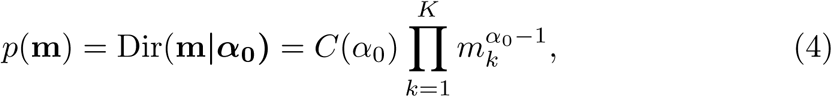

where *C*(*α*_0_) is the normalizing constant for the Dirichlet distribution.

We also take group parameters ***µ*** and **T** as random variables, which allows us to evaluate their posterior distribution given data. We introduce conjugate priors for these variables, a Gaussian-Gamma prior in which the distribution over ***µ***_*k*_ depends on **T**_*k*_:

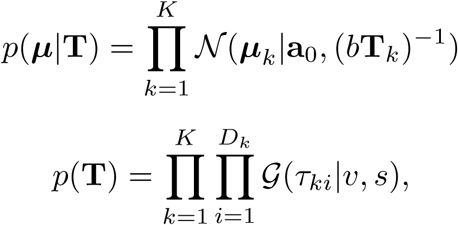

where *𝒢* (.) denotes Gamma distribution. Here, *τ*_*ki*_ is the *i*th diagonal element of **T**_*k*_. Assuming that ***τ*** _*k*_ is a vector containing *τ*_*ki*_, by defining **T**_*k*_ = diag(***τ*** _*k*_), in which diag(.) is an operator outputting a diagonal matrix with elements given by ***τ*** _*k*_, we can write these two equations in a compact form:

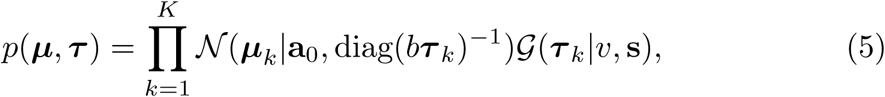

where we have defined:

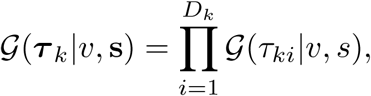

in which 𝓋 is a scalar and **s** is a vector with *D*_*k*_ elements all equal to *s*. The full probabilistic model is given by,

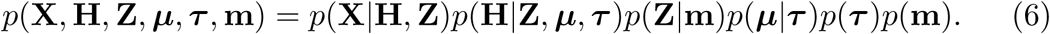

### A.2 Variational inference

The task of Bayesian inference is to compute the posterior probabilities of latent variables given data, *p*(**H**, **Z**, ***µ***, ***τ***, **m**|**X**). Since the inference is intractable for the probabilistic model outlined in section A.1, we employ variational inference to compute approximate posteriors. We take a so-called mean-field approach [15] by assuming that the posterior is partially factorized as follows:

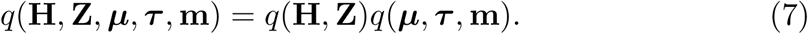

Note that we force no factorization in the posterior between latent variables, **Z** and **H**. Using a quadratic approximation of the conditional posterior, *q*(**H**|**Z**), we prove in Appendix B that these posteriors are given by,

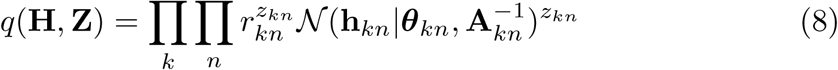

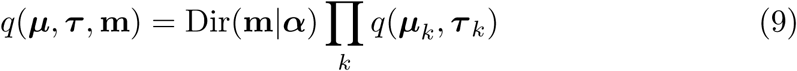

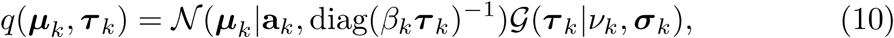

where *𝓋*_*k*_ and *β*_*k*_ are scalars and *σ*_*k*_ is a vector with the same size as ***τ*** _*k*_. In the next section, we provide the HBI algorithm, which iteratively updates the parameters of these distributions, *r*_*kn*_, *θ*_*kn*_, **A**_*kn*_, ***α***, **a**_*k*_, *𝓋*_*k*_, *β*_*k*_, and *σ*_*k*_.

### A.3 HBI algorithm

After initializing the individual parameter estimates, *θ*_*kn*_ and **A**_*kn*_ and responsi-bilities *r*_*kn*_ for all subjects and models, as well as setting prior parameters **a**_0_, *b, s*, 𝓋 and *α*_0_ (see A.6 for a simple and intuitive way for initializing and setting prior parameters), the HBI algorithm performs these steps:

- 1. Calculate the summary statistics:

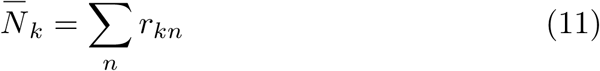

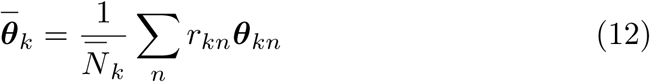

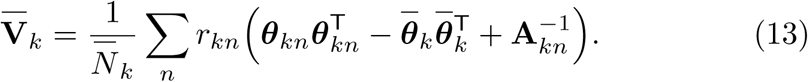
- 2. Update parameters of *q*(***µ***, ***τ***, **m**) for all models:

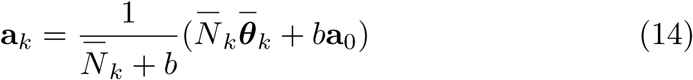

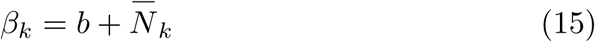

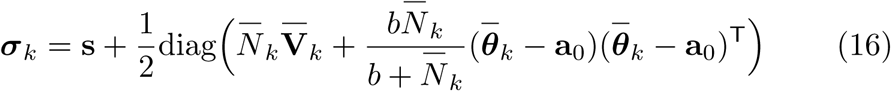

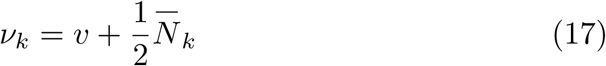

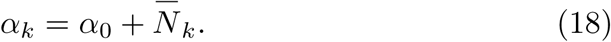
- 3. Update the individual posterior parameters *θ*_*kn*_, **A**_*kn*_ and *f*_*kn*_, by obtaining a quadratic approximation of the function, *l*_*kn*_(h), with respect to **h**:

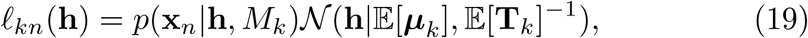

where 𝔼 [***µ***_*k*_] = **a**_*k*_ and 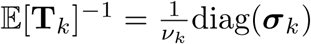 diag. This approximation can be written as

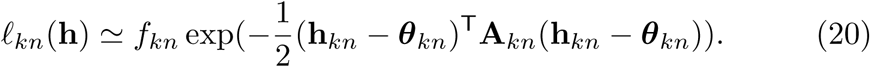 Note that any quadratic approximation can be used here. For example, using a Laplace quadratic approximation (which is a very common approximation for analyzing behavioral and neural data [19-22]), *θ*_*kn*_, **A**_*kn*_ and *f*_*kn*_ are given by the mode, Hessian of log *f*_*kn*_ and the maximum value of *l*_*kn*_, respectively:

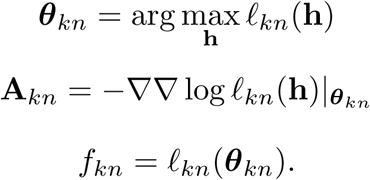
- 4. Update responsibilities,

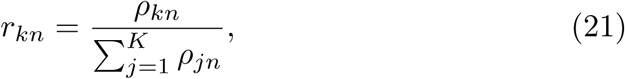

where

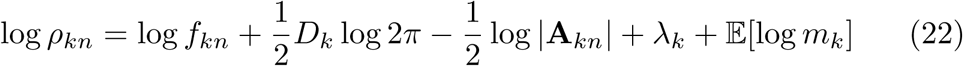

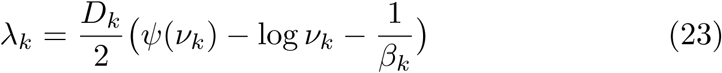

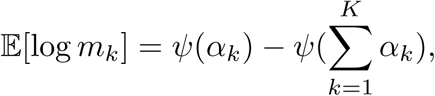

in which ψ (.) is the digamma function.
- 5. Terminate if stopping criteria are met, otherwise go to 1.

### A.4 Statistical tests for group parameters

An important goal of computational modeling studies is to compute the distribution of parameters given data across the whole population. From a Bayesian viewpoint, this is given by the marginal posterior over the mean of group parameters, ***µ***_*k*_, which reads

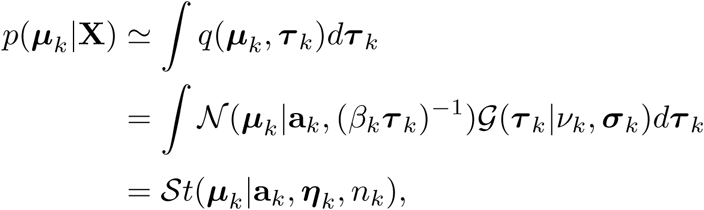

where 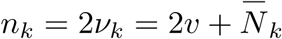 is the number of degrees of freedom of the Student distribution and 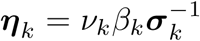 is the inverse-scale parameter. Therefore, the random variable 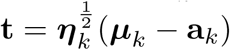 takes a form of standard Student distributio with *n*_*k*_ degrees of freedom. By defining 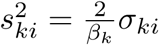, in which 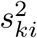 corresponds to empirical variance (c.f. equation (16)), we can write this result in an intuitive form,

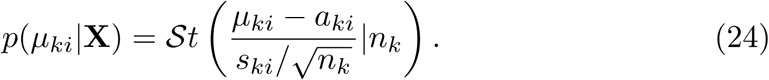

Noting the similarity between 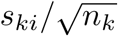 and the standard error of the mean, we called 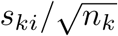 the hierarchical error.

### A.5 Predictive distribution for a new subject

In many situations, researchers are interested to fit a new dataset to a particular model and find corresponding parameters. In Bayesian statistics, this is called the predictive distribution and it is given by marginalizing over group parameters. Suppose that **x**^*\**^ and 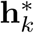 denote the new dataset and its corresponding parameters for model *k*. The marginal distribution 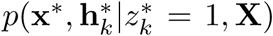 is the predictive distribution given the observed dataset **X** assuming that the new data is generated by the *k*th model. This distribution is given by:

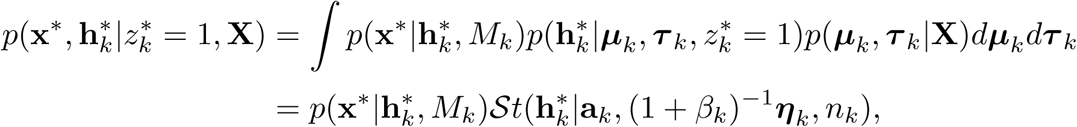

where *η*_*k*_ and *n*_*k*_ have been defined in the previous section. This distribution can also be written in terms of standard Student distribution with *n*_*k*_ degrees of freedom. Furthermore, if we assume that *b* = 2𝓋, which is a reasonable assumption (see the next section), this distribution is given by

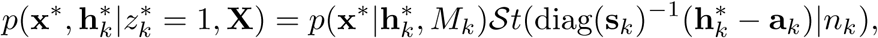

where s_*k*_ is a vector of corresponding empirical deviance parameters, defined in the previous section. Using this joint distribution, one can use sampling methods to obtain the posterior over parameters, 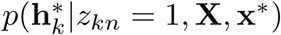, or to obtain the maximum-a-posteriori parameters,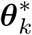, given by

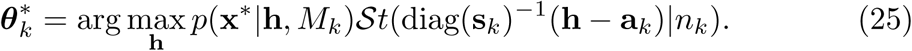

Note that for many degrees of freedom due to large values of 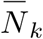, the Student distribution tends to a Gaussian with mean a_*k*_ and deviance matrix diag(s_*k*_). However, small 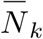 lead to a small number of degrees of freedom and heavier tailed distributions than Gaussians, which are more robust against outliers.

### A.6 Parameters, initialization and convergence criteria

As the mean-field variational inference is an iterative framework, it also depends on the initialization of the parameters. In this section, we provide priors that do not bias the final solution and also provide some intuitive criteria for the initialization.

We initialize the parameters *θ*_*kn*_ and_*kn*_ by fitting all models separately to all participants (with some initial Gaussian prior), i.e., assuming as if *z*_*kn*_ = 1. These values are then used to calculate summary statistics according to equations (11-13).

Furthermore, we need to define prior parameters. The free parameter *α*_0_ indicates prior frequency of each model. We take uninformative priors on frequency of models, which is given by *α*_0_= 1 for all models. The prior mean, a_*0k*_, is assumed to be zero. Given equation (15), we see that *b* can be interpreted as the effective number of prior samples associated with models. Also, given equation (17), *𝓋* could be interpreted as the half of the effective number of prior samples associated with models. Assuming that the priors account for one sample, which is a common assumption in Bayesian statistics, we take *b* = 1 and 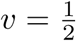. Finally, since *s* has always an additive effect on *σ*_*k*_ according to equation (16), we assume a small positive value for *s*, allowing that *σ*_*k*_ to be driven dominantly by data. In all our analyses, we assumed *s* = 0.01.

Finally, the algorithm presented in A.3 requires stopping criteria. In our analyses, we terminated the algorithm if the change in normalized value of parameters between two consecutive iterations, *j -* 1 and *j*, defined as

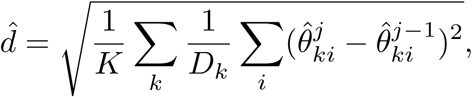

was smaller than 0.01. Here,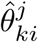: is defined according to summary statistics of parameters on the *j*th iteration

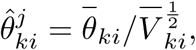

where *θ*_*ki*_ and 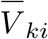 are the *i*th element of 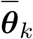 and 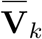 defined in (12-13), respectively. In our analyses, we also set 50 as the maximum number of iterations, although almost always the algorithm stopped before hitting this number.

### A.7 Exceedance probability

Using the posterior over **m**, one can also derive the so-called exceedance probability and protected exceedance probability, as defined in previous works [13,18]. We reproduce the equations here for completeness.

The exceedance probability of *k*th model, *φ*_*k*_, is defined as the probability that model *M*_*k*_ is more likely than any other model in the model-space and it is given by

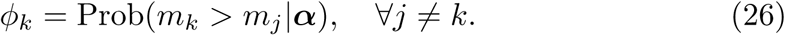

Computing protected exceedance probabilities, as defined in [18], also requires to run the HBI under the (prior) null hypothesis, *H*_0_, that there is no difference between models (i.e. *α*_0_ → *∞*). The alternative hypothesis, *H*_1_, is the original case, in which *α*_0_ = 1. If we define *L* and *L*_0_ as the log-likelihood (actually the variational lower bound as its approximation) of all data given the model-space under *H*_1_ and *H*_0_, respectively, then the protected exceedance probability of *k*th model,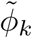, is defined as:

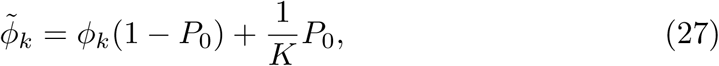

where

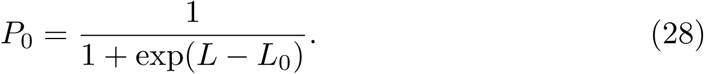

## Appendix B

In this appendix, we provide the proof of the results given in sections A.2 and A.3. The proof is given in three parts by obtaining 1) the functional form of *q*(**H**, **Z**); 2) the posterior *q*(***µ***, ***τ***, **m**) and corresponding update equations; and 3) the update equations for the posterior over latent variables, *q*(**H**, **Z**).

### B.1 The functional form of the posterior over H and Z

Let us first consider the derivation of the functional form for the factor *q*(**H**, **Z**). According to standard results in variational inference [15,16], the log of this factor is given by:

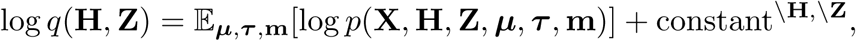

where the constant term denotes all the terms independent of the corresponding variables. Note that the expectation is taken with respect to the current estimates of *q*(***µ***, ***τ***, **m**). By using equation (6) and absorbing all the terms which are independent of **H** and **Z** into the additive constant, we have:

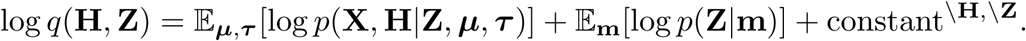

Substituting the two conditional distribution on the right-hand side using equations (1-3), we have:

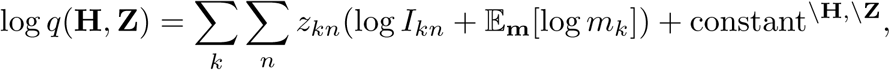

where,

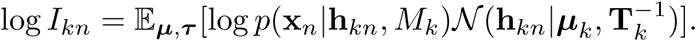

Note that we have defined **T**_*k*_ = diag(***τ*** _*k*_). We assume that there is a quadratic approximation of *I*_*kn*_ with respect to **h**_*kn*_,

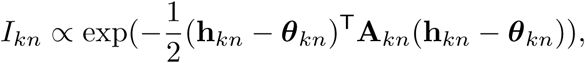

which gives,

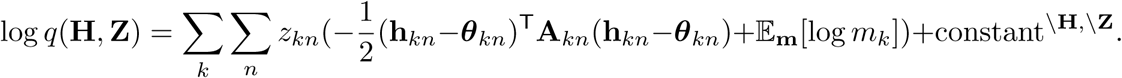

Since log *q*(**H**|**Z**) = log *q*(**H**, **Z**) - log *q*(**Z**), we can read off terms involving **H** in log *q*(**H**, **Z**) to obtain log *q*(**H**|**Z**):

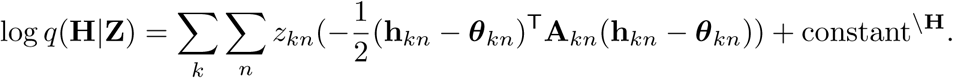

Requiring that this distribution should be normalized, we obtain:

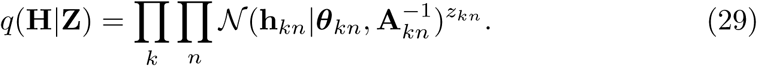

Subtracting log *q*(**H**|**Z**) from log *q*(**H**, **Z**) cancels out the quadratic component and yields log *q*(**Z**), which is a linear function with respect to *z*_*kn*_. Therefore, we have:

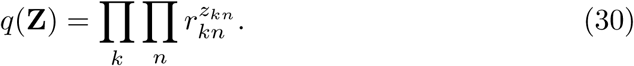

The functional form of *q*(**H**, **Z**) is then given by,

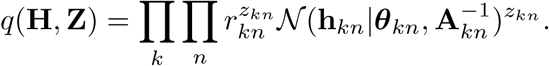

Here our goal was to obtain the functional form of the posterior over latent variables. We will obtain values of *r*_*kn*_, *θ*_*kn*_ and **A**_*kn*_ in section B.3.

### B.2 The posterior over *µ, τ* and m

We continue with obtaining the functional form and update equations for the other variational factor *q*(***µ***, ***τ***, **m**). The posterior of **m** is independent from the posterior over ***µ*** and ***τ*** because the log-posterior decomposes into the terms that only depend on **m** and terms that only depend on ***µ*** and ***τ***:

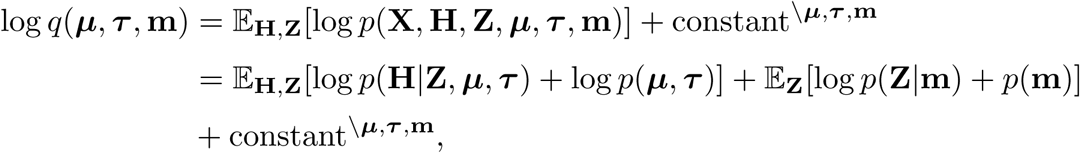

where we have used equation (6). This implies that the variational posterior *q*(***µ***, **T**, **m**) factorizes to give *q*(***µ***, **T**)*q*(**m**). Thus, the posterior over ***µ*** and ***τ*** is given by:

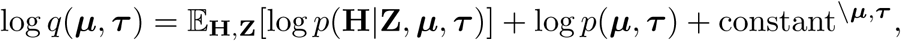

in which we absorbed any terms independent of ***µ***, ***τ*** into the additive constant. Substituting for the distributions on the right-hand side, we have:

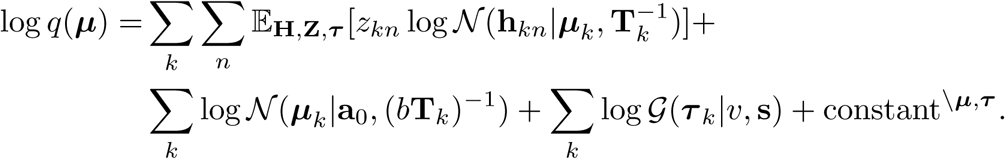

Note that we have defined **T**_*k*_ = diag(***τ*** _*k*_). Using equation (B.1) and by absorbing terms independent of ***µ***_*k*_ and ***τ*** _*k*_ into the additive constant, we have:

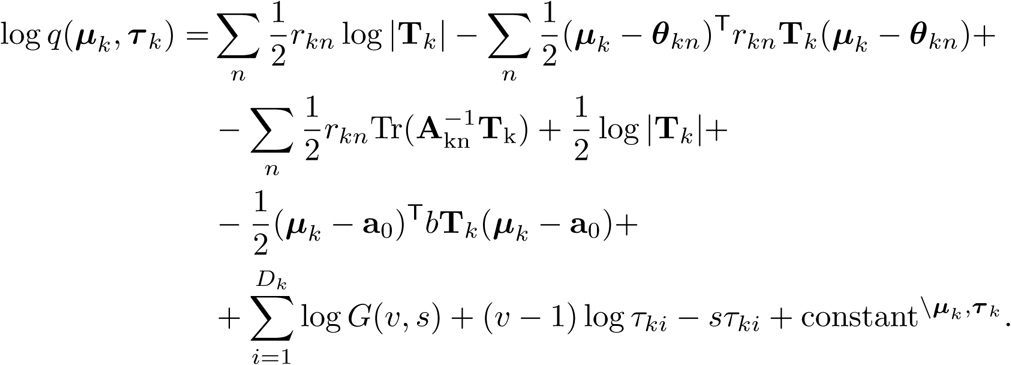

As the right-hand side is quadratic with respect to ***µ***_*k*_, the posterior over ***µ***_*k*_ also takes the form of a Gaussian with a variance depending on ***τ*** _*k*_:

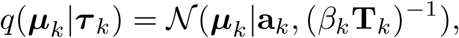

where

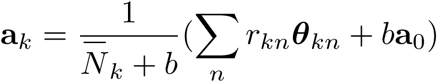

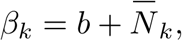

and *N*_*k*_ is given by:

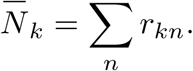

By subtracting log *q*(***µ***_*k*_|***τ*** _*k*_) from log *q*(***µ***_*k*_, ***τ*** _*k*_), we obtain the posterior over ***τ*** _*k*_:

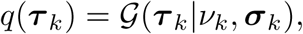

where

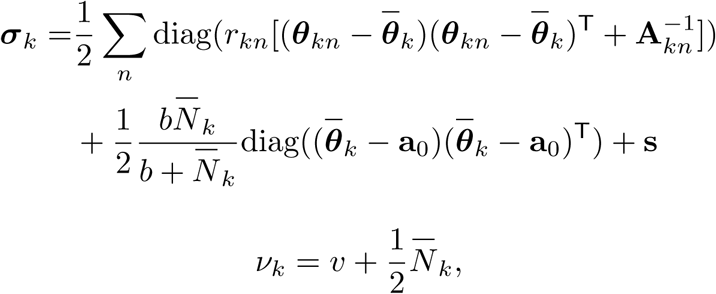

and 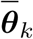 is given by:

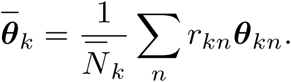

Finally, we consider the factor *q*(**m**):

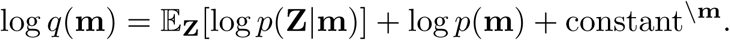

Substituting for the two distributions on the right-hand side, we have

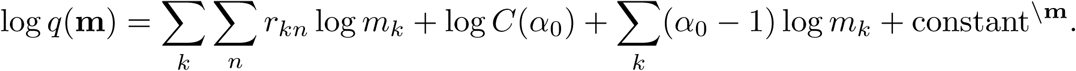

Therefore *q*(**m**) takes the form of Dirichlet distribution:

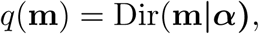

where *α* has components *α*_*k*_ given by,

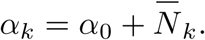

### B.3 The posterior over H and Z

We have already seen in section B.1 that *q*(**H**, **Z**) could be written,

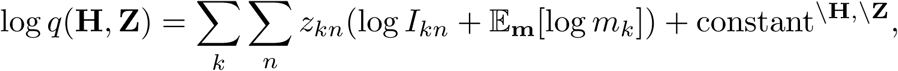

where

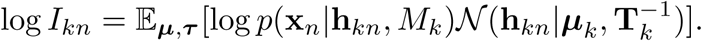

Since we have already obtained *q*(***µ***, ***τ***), we can now compute *I*_*kn*_:

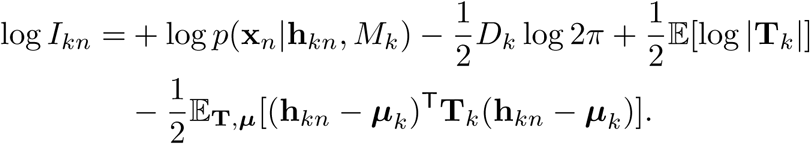

By using equation (9), log *I*_*kn*_ is given by:

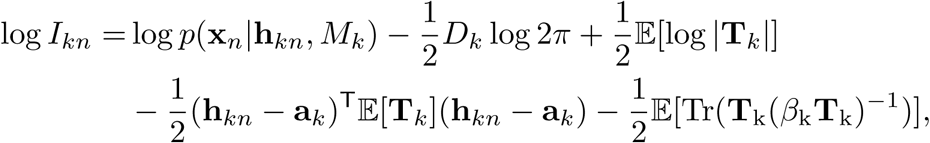

which can be written in the form,

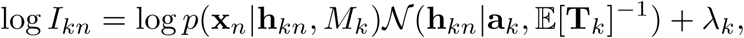

where *λ*_*k*_ is independent of **h**_*kn*_ and is given by

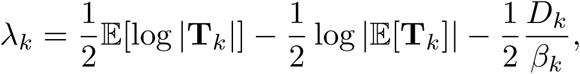

Substituting the moments of **T**_*k*_ = diag(***τ*** _*k*_) with their values under *q*(***τ*** _*k*_),

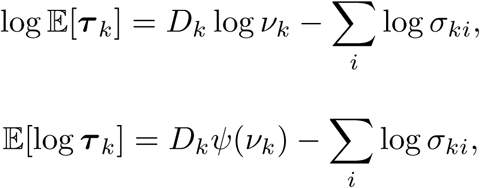

gives equation (23).

Now, we make a quadratic approximation of *p*(**x**_*n*_|**h**_*kn*_, *M*_*k*_) 𝒩 (**h**_*kn*_|a_*k*_, 𝔼 [**T**_*k*_]^-1^) with respect to h_*kn*_ (for example using Laplace approximation or any other method):

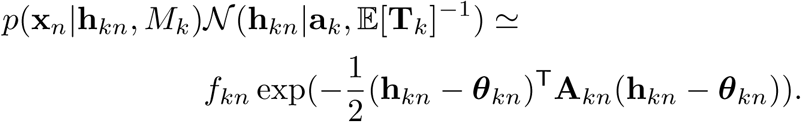

Substituting this approximation into log *I*_*kn*_, we obtain

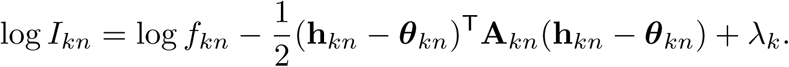

Therefore, we have:

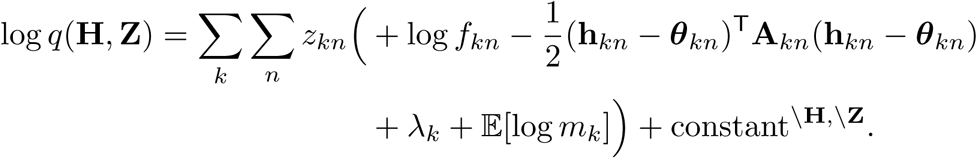

Subtracting this equation from log *q*(**H**|*Z*) given by the log of equation (29), we have:

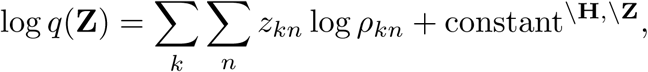

where

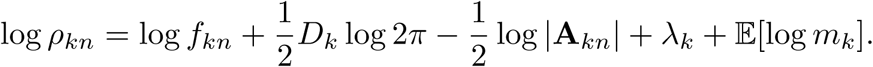

Requiring that *q*(**Z**) be normalized, we obtain equation (30), where

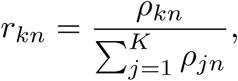

which completes the proof.

